# Inter-Subject Alignment of MEG Datasets in a Common Representational Space

**DOI:** 10.1101/096040

**Authors:** Qiong Zhang, Jelmer P. Borst, Robert E. Kass, John R. Anderson

## Abstract

Pooling neural imaging data across subjects requires aligning recordings from different subjects. In magnetoencephalography (MEG) recordings, sensors across subjects are poorly correlated both because of differences in the exact location of the sensors, and structural and functional differences in the brains. It is possible to achieve alignment by assuming that the same regions of different brains correspond across subjects. However, this relies on both the assumption that brain anatomy and function are well correlated, and the strong assumptions that go into solving the underdetermined inverse problem given the high dimensional source space. In this paper, we investigated an alternative method that bypasses source-localization. Instead, it analyzes the sensor recordings themselves and aligns their temporal signatures across subjects. We used a multivariate approach, multi-set canonical correlation analysis (M-CCA), to transform individual subject data to a low dimensional common representational space. We evaluated the robustness of this approach over a synthetic dataset, by examining the effect of different factors that add to the noise and individual differences in the data. On a MEG dataset, we demonstrated that M-CCA performs better than a method that assumes perfect sensor correspondence and a method that applies source localization. Lastly, we described how the standard M-CCA algorithm could be further improved with a regularization term that incorporates spatial sensor information.

## 1. Introduction

In neuroimaging studies, data are frequently combined from many subjects to provide results that represent a central tendency across a population. To achieve this, it is necessary to find an alignment between brain activities recorded from different subjects. In MEG recordings, corresponding sensors across subjects can be poorly correlated both because of differences in the exact location of the sensors^1^ (i.e. different head position in the helmet), and structural and functional differences in the brains.

It is possible to achieve alignment by assuming that the same regions of different brains correspond across subjects. Corresponding brain regions are found by first localizing brain sources for each subject. The most commonly used source localization method computes the minimum-norm current estimates (MNE) in an inverse modeling approach (Gramfort et al., 2014). In a separate second step, typically using individual MRIs, the sources for individual subjects are morphed onto sources in a common ‘average’ brain^2^. The resulting morphed sources are considered to correspond from subject to subject. The validity of this approach depends both on the assumption that brain anatomy and function are well aligned and on the strong assumptions that go into solving the inverse problem of source localization. These steps and assumptions are satisfactory as long as the localization errors combined with the distortions in morphing are small relative to the effects being investigated.

In this paper, we proposed the use of multi-set canonical correlation analysis (M-CCA) to transform individual subject data to a low dimensional common representational space where different subjects align. The transformation is obtained by maximizing the consistency of different subjects’ data at corresponding time points^3^ (Kettenring, 1971). This way, our approach utilizes the rich temporal information that MEG sensor data offers and circumvents the need to find anatomical correspondence across subjects. This gives many advantages. Firstly, M-CCA does not rely on any assumptions that go into solving the source localization problem. Secondly, M-CCA focuses on capturing distributed patterns of activity that have functional significance, and establishes correspondence across subjects based on these patterns. Therefore, M-CCA does not rely on the assumption that brain anatomy and functions are well correlated. It simultaneously deals with three factors that contribute to subject misalignment in sensor data: 1) the structural differences of the brains; 2) the sensor location differences during MEG recordings across subjects; and 3) the different structure-function mappings in different subjects.

The output of the M-CCA analysis can be used in multiple ways. For example, one might want to find the time windows over which the differences between conditions are decoded in the neural signal (Norman et al., 2006). M-CCA allows one to go from intra-subject classification to inter-subject classification where one group of subjects can be used to predict data in a new subject. M-CCA may also be used to build a group model that studies the sequence of temporal mental stages in a task (Anderson et al., 2016). Although temporal data from individual subjects are too sparse to be analyzed separately, once the data from all subjects are aligned, they can be combined to improve model parameter estimation. Once important time windows of activity modulation have been identified in such a group model, it is possible to go back to individual subjects and observe the corresponding localized signals.

M-CCA has been applied previously in fMRI studies. Rustandi et al. (2009) applied the M-CCA method to achieve successful prediction across multiple fMRI subjects. Li et al. (2012) used M-CCA to integrate multiple-subject datasets in a visuomotor fMRI study, where meaningful CCA components were recovered with high inter-subject consistency. Correa et al. (2010) reviewed a wide range of neuroimaging applications that would potentially benefit from M-CCA. This not only includes a group fMRI analysis pooling multiple subjects, but also fusion of data from different neuroimaging modalities (i.e. fMRI, sMRI and EEG). MEG is very different from fMRI in terms of temporal resolution and the way the underlying neural signal is generated. To align MEG data, in this paper we first proposed the pipeline of applying M-CCA to MEG data. Second, we evaluated the suitability of the method with a synthetic dataset that is realistic to MEG recordings, which models: 1) the generation of the neural signal in the brain sources; 2) the mapping from activity at the brain sources to activity at the sensors; and 3) the noise added to the sensors during the measurement. We explored different factors that could add to the noisiness of the data, including increased levels of noise added to the sensors, and continuous head movement during the recordings that can further blur the sensor measurement (Stolk et al., 2013; Uutela et al., 2001). We also explored different factors that could add to the individual differences in the data, including variation in brain patterns across subjects, different noise conditions across subjects, and unique functional brain responses in each subject. M-CCA performed robustly in the presence of these factors.

Finally, we demonstrated on a real MEG dataset that M-CCA performed better than a method that assumes perfect sensor correspondence and a method that applies source localization. We also showed how the standard M-CCA algorithm could be further modified to take into account the similarity of M-CCA sensor mappings across different subjects. A documented package has been made available to pre-process data and apply M-CCA to combine data from different MEG subjects (https://github.com/21zhangqiong/MEG_Alignment).

## 2. Materials and Methods

### 2.1. MEG Experiment

Twenty individuals from the Carnegie Mellon University community completed the experiment, which was originally reported in Borst et al. (2016). Two subjects were excluded from analysis (one fell asleep and one performed subpar). All were right-handed and none reported a history of neurological impairment. The experiment consisted of two phases: a training phase in which subjects learned word pairs and a test phase in which subjects completed an associative recognition task. The test phase was scheduled the day after the training phase and took place in the MEG scanner. During the test phase subjects had to distinguish between targets (learned word pairs) from foils (alternative pairings of the learned words). Subjects were instructed to respond quickly and accurately. There were four binary experimental factors: probe type (targets or foils), word length (short: with 4 or 5 letters; long: with 7 or 8 letters), associative fan (one: each word in the pair appears only in that pair; two: each word in the pair appears in two pairs), and response hand (left or right). Subjects completed a total of 14 blocks (7 with left-handed responses, 7 right-handed), with 64 trials per block. MEG data were recorded with a 306-channel Elekta Neuromag (Elekta Oy) whole-head scanner, which was digitized at 1 kHz. A band-pass filter (0.5−50 Hz) was applied using FieldTrip toolbox (Oostenveld et al., 2011) before the data was down-sampled to 100 Hz ^4^. As a pre-processing step to compensate for head movement across different sessions on an individual subject, sensor data were realigned to average head positions using MNE-based interpolation (Hämäläinen & Ilmoniemi, 1994). Alternative methods to correct for head movement exist, including the signal space separation (SSS) method (Nenonen et al., 2012) and an extension of SSS-based method that takes care of more challenging cases of head movement across different age groups (Larson & Taulu, 2016). More details of the experiment can be found in the original report of this MEG dataset (Borst et al., 2016).

### 2.2. Alignment by Correspondence of Brain Sources

We compare M-CCA with two other methods. The first method assumes that the same sensors correspond across subjects. The second method is to perform source localization first, and then assume that the same sources correspond across subjects. In MEG recordings, the measured magnetic signal does not directly indicate the location and magnitude of cortical currents, which can be found by projecting the sensor data onto the cortical surface with minimum norm estimates (MNE). The MNE method attempts to find the distribution of currents on the cortical surface with the minimum overall power that can explain the MEG sensor data (Gramfort et al., 2014). This is done by first constructing 3D cortical surface models from the subjects’ structural MRIs using FreeSurfer, which are then manually co-registered with the MEG data (Dale et al., 1999; Fischl, 2012). A linear inverse operator is used to project sensor data onto the source dipoles placed on the cortical surface. These source estimates are then morphed onto the standard MNI brain using MNE’s surface-based normalization procedure. Source estimates on the standard MNI brain are thought to correspond across subjects. More details for obtaining source localization with MNE over the current MEG dataset can be found in the original report (Borst et al., 2016).

### 2.3. Alignment in the Common Representational Space

This section outlines our proposed approach to align MEG subjects. M-CCA is used to find the optimal transformation for each subject from the activity of 306 sensors to a common representational space. Inter-subject correlations of the transformed data are maximized across subjects in the common representational space.

#### 2.3.1. Application of M-CCA to MEG Data

This section describes the pipeline to apply M-CCA to MEG datasets. We start by pre-processing the sensor data for each subject. To overcome noise in the sensor data of individual trials, multiple trials are averaged to obtain a highly reliable representation of the change in sensor activity, which is similar to obtaining event-related potential waveforms in the EEG literature (Picton et al., 2000). When trials have a fixed trial length, this averaging procedure is straightforward. However, in the current experiment, trials are variable in their durations, as the responses made by participants are self-paced. In order to align time samples across trials before averaging, we consider only the samples from the first half second (50 samples given the sampling rate) and the last half second (another 50 samples) of a trial in our application of M-CCA. Neural signals of the same cognitive event have less trial-to-trial variability when they are closely locked to the stimulus/response. When they are further away from the stimulus/response, they are less reliably recovered in the average signal. The process of averaging is repeated for each condition, as we potentially have different latent components for different conditions after averaging. With four binary factors (probe type, word length, associative fan, response hand), we have 16 conditions. Essentially, over an MEG dataset, the transformation of subject data in M-CCA is obtained by maximizing the consistency of different subjects both in response to different temporal points in a trial, and in response to different experimental conditions. With 100 samples per condition after averaging over trials, we have a 1600 × 306 matrix *S_k_* from 306 sensors for each of 18 subjects as the initial input to the M-CCA procedure (Figure 1). This format of input data is very different to that of fMRI data where temporal resolution within a single trial is limited. To have enough temporal information to align multiple datasets, M-CCA is typically applied to fMRI data either using a long stream of continuous scans or a series of trials where there is unique temporal information for each trial (Li et al., 2012; Rustandi et al., 2009). The finer temporal resolution in MEG opens up opportunities to use M-CCA to align subjects in a wider range of experimental tasks (e.g. repeated-trial design).

**Figure 1:**
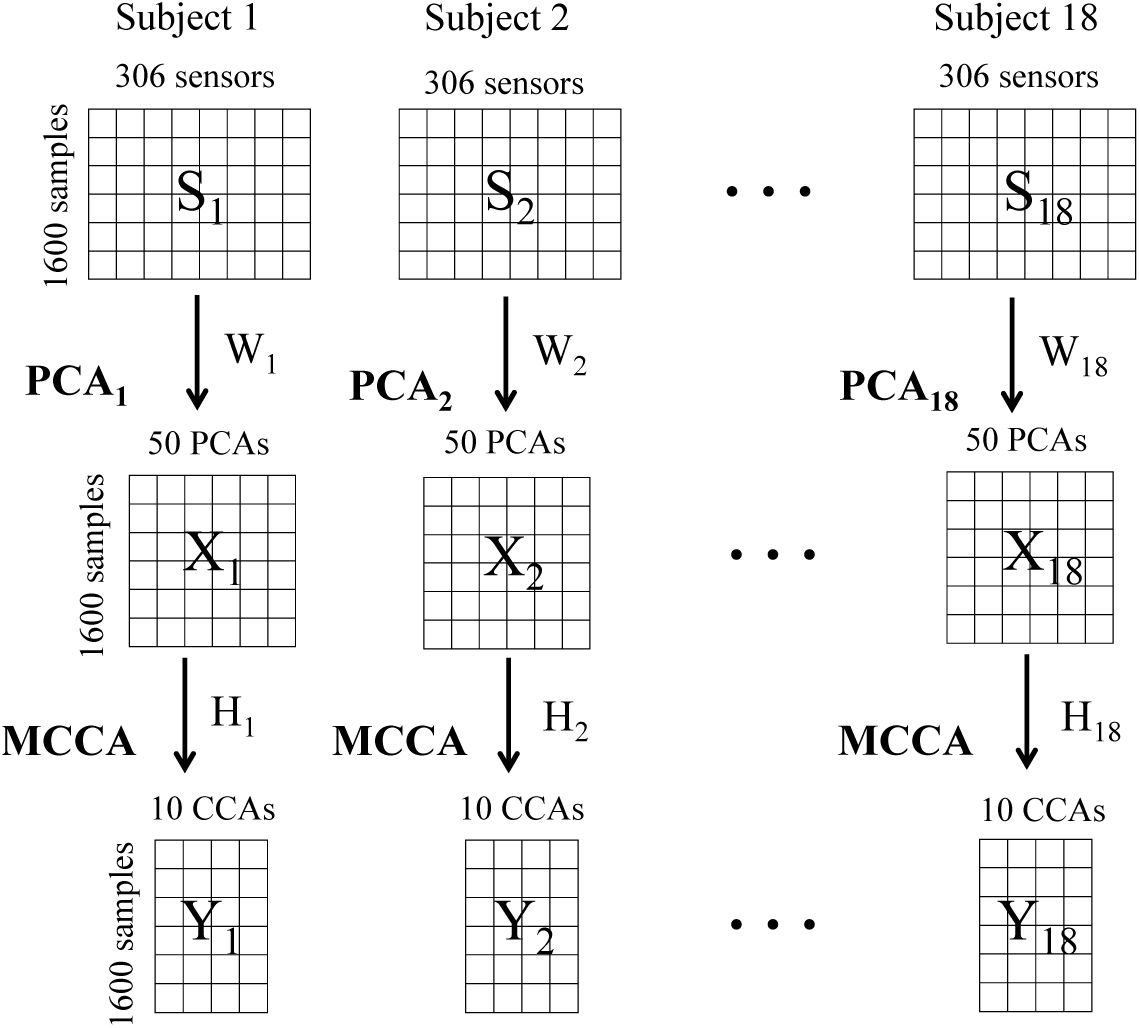
This figure illustrates application of M-CCA to 18 subjects. *S_k_* is the averaged data (across all trials for each condition) from 306 sensors for subject *k*, each with 1600 time points; *X_k_* has 50 PCA components for subject *k*, each with 1600 time points; *Y_k_* has 10 CCA components, each with 1600 time points; *W*_1_*,W*_2_*,…, W*_18_ are PCA weights obtained for each subject independently; and *H*_1_*, H*_2_*,…, H*_18_ are CCA weights obtained jointly from all subjects by maximizing all of the inter-subject correlations.

To reduce dimensionality and remove subject-specific noise, the next step after obtaining *S_k_* is to perform spatial PCA. M-CCA is then applied to the top 50 PCA components from each subject instead of directly to the sensor data *S_k_*. This results in 18 matrices of dimension 1600 × 50, which are the inputs *X_k_* to the M-CCA analysis for subjects *k* = 1, 2,…, 18. As is illustrated in Figure 1, *W_k_* are the PCA weights for subject *k* which are obtained independently for each subject. *H_k_* are the CCA weights for subject *k* which are obtained jointly from all subjects resulting in common CCA dimensions *Y_k_* = *X_k_H_k_*. Subject data do not align in either the sensor space or the PCA space, with *S_k_* and *X_k_* processed for each subject independently. Rather, subject data align in the common representational space after M-CCA, with *Y_k_* maximally correlated across subjects. The selection of number of PCA components and CCA components will be discussed in the results section.

#### 2.3.2. M-CCA

This section discusses the mathematical details of how *H_k_* for subjects *k* = 1, 2*,…,* 18 are obtained, so that after the transformation *Y_k_* = *X_k_H_k_*, the new representation of data *Y_k_* is more correlated across subjects than *X_k_* is.

We first illustrate the simplest case where we look for correspondence over datasets from two subjects instead of many subjects. Let 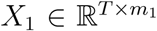 and 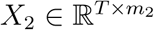 be PCA components from two subjects, with the same number of time points *T*, and PCA dimensions *m*_1_ and *m*_2_, respectively (*T* = 1600, *m*_1_ = *m*_2_ = 50 in our case). Each PCA component stored in *X*_1_ and *X*_2_ has mean 0. The objective in canonical correlation analysis (CCA) is to find two vectors 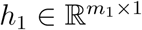 and 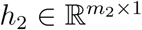 such that after the projection *y*_1_ = *X*_1_*h*_1_ and *y*_2_ = *X*_2_*h*_2_, *y*_1_ and *y*_2_ are maximally correlated. This is equivalent to:

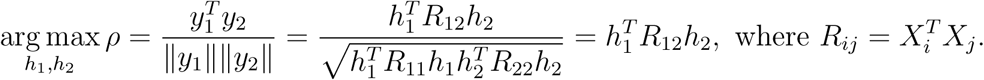

There are *N* solutions, 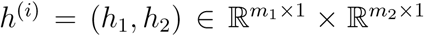, obtained collectively in a generalized eigenvalue problem with *i* = 1*,…, N*, subject to the constraints 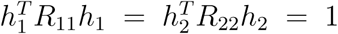 (Borga, 1998). This results in *N* dimensions (each referred as a CCA component) in the common representational space with the transformed data 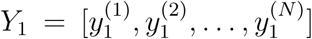 and 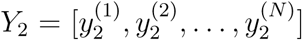. The value of *N* does not exceed the smaller of *m*_1_ and *m*_2_. The resulting CCA components in the common representational space are ranked in a decreasing order of between-subject correlations. The earlier CCA components are the more important ones and the later components can be removed. In other words, canonical correlation analysis finds the shared low-dimensional representation of data from different subjects.

M-CCA is an extension of CCA which considers more than 2 subjects. The objective is similar to before, but now it needs to maximize the correlations between every pair of subjects (i.e. inter-subject correlations) simultaneously. Let 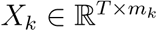 with *k* = 1*,…, M* be datasets from *M* subjects (*M >* 2), each with mean 0 for all columns. The objective in M-CCA is to find *M* vectors 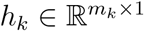, where *k* = 1*,…, M*, such that after the projection *y_k_* = *X_k_h_k_*, the canonical variates *y_k_* are maximally pairwise-correlated. The objective function to maximize is formulated as:

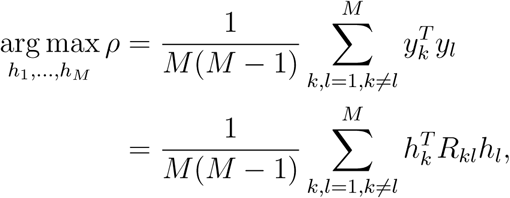

where 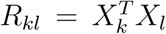 and 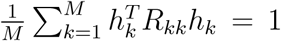. The solution is given by solving a generalized eigenvalue problem (Vía et al., 2007). This formulation is not an exact maximization but an approximation of the pairwise correlations, given the complexity of the problem when *M* > 2. It is equivalent to the Maximum Variance (MAXVAR) generalization of CCA proposed by Kettenring (1971). See the proof of this equivalence in (Vía et al., 2005). Other ways of formulating the objective function in M-CCA yield similar results (Li et al., 2009).

#### 2.3.3. M-CCA with a Regularization Term

To further improve the application of M-CCA on subject alignment, we also take into consideration the spatial sensor information across different subjects. Projection weights that map sensor data to a given CCA component should be similar across different subjects, given that the misalignment potentially results from small spatial shifts either during sensor recordings or from anatomical variation across subjects. We enhance this similarity (captured as the correlations of sensor weight maps across subjects) by adding a regularization term in the M-CCA algorithm. This term is useful in situations where there are multiple projection weights that give rise to similar results at a given CCA dimension due to the highly correlated sensor activities. Adding a regularization term makes sensor-to-CCA projection weights similar across subjects while simultaneously maximizing the inter-subject correlations of the transformed data. This leads to more interpretable and unique projection weights, and potentially improves the resulting CCA dimensions under certain scenarios.

Let 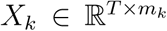, where *k* = 1*,…, M*, be PCA components obtained independently for each of the *M* subjects (*M >* 2), *T* be the number of time points, and *m_k_* be the number of PCA components. Let 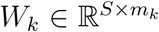, where *k* = 1*,…, M*, be the subject-specific PCA weights (sensor-to-PCA projection) from *M* subjects, and *S* be the number of MEG sensors. The modified M-CCA is formulated as:

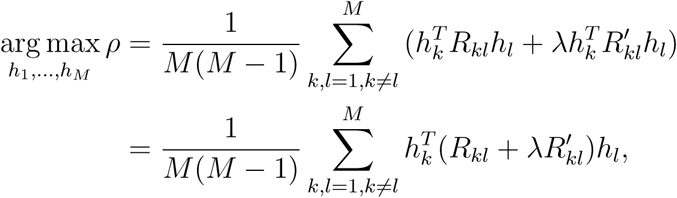

where 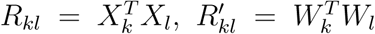, and 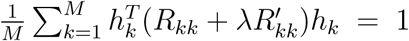 Essentially, the inter-subject correlations of the transformed data *X_k_h_k_* are over *T* time points, whereas the inter-subject correlations of the sensor-to-CCA mapping *W_k_h_k_* are over *S* sensors. This explains why the maximization of the two components in the objective function takes a very similar format, with only the weights differing. One can vary the emphasis given to the regularization term via the scalor *λ*. *λ* = 0 corresponds to the M-CCA algorithm without regularization, and very large values of *λ* correspond to using the same weight maps for all subjects. The solution to the M-CCA problem with regularization can be obtained in the same way as the standard M-CCA by solving a generalized eigenvalue problem.

### 2.4. Generation of MEG Synthetic Datasets

To test the applicability of M-CCA on MEG datasets, we generate synthetic datasets that are realistic to MEG recordings. This includes modeling the generation of the neural signal in the brain sources, the mapping from currents at the brain sources to activities measured at the sensors, and the noise added to the sensor during the measurement. The procedure is done separately when simulating each new subject. Therefore, we can model subject misalignment in sensor data explicitly, by adding individual differences in: 1) structure of the brain; 2) sensor locations; 3) structure-function mappings; and 4) noise level during the recording. The first two factors of individual differences are taken into account by using the forward models obtained directly from the 18 experimental subjects using the MNE software. Brain sources are modeled as current dipoles (Scherg & Von Cramon, 1985; Hamalainen & Sarvas, 1989), and a forward model specifies how currents at the dipoles map to activities measured at the sensors. This is calculated for each subject separately using a boundary element model (BEM) based on individual head geometry and the configuration of the MEG sensors (Oostendorp & van Oosterom, 1989). Head geometry is used to approximate the distribution of magnetic and electrical fields (Gramfort et al., 2010; Mosher et al., 1999). Sensor properties includes locations, orientations and coil geometries (i.e. magnetometer, axial and planar gradiometers). As MEG is mainly sensitive to electric currents in the pyramidal cells (Hämäläinen et al., 1993), we use forward models that constrain dipole currents along the normal direction of cortical surfaces. The last two factors of individual differences, structure-function mappings and noise levels, are what we can vary in the synthetic datasets to examine the robustness of the M-CCA procedure. Structure-function mappings refer to the possibility that even if subjects have the same brain structures, different brain regions can be recruited to perform the same task.

Generation of single-trial sensor data for each subject *k* can be described as:

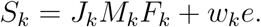

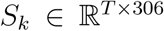 represents single-trial data of length *T* recorded at 306 sensors. 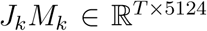 represents the source data of the 5124 dipoles, with 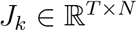 being *N* independently generated temporal components (i.e. underlying brain dynamics), and 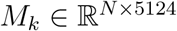 being a spatial mask consisting of 1s and 0s that specifies if a dipole is involved in a particular temporal component. *F_k_* is the forward model for subject *k* that maps dipole data to sensor data. *e* is the environmental noise added to the sensors, which is generated at each time point *t* as

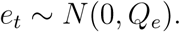

The noise covariance *Q_e_* is obtained from empty room recordings. *w_k_* models the individual variability in the noise level given that different subjects are recorded during different recording sessions. Signal-to-noise ratio (SNR) measures the power of brain signal *J_k_M_k_F_k_* divided by the power of sensor noise *w_k_e*.

Further, we have 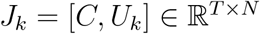, where *N* = *n* + *m*. 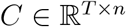 represent *n* temporal components that are shared across all subjects, which are set to be the top 10 CCA components obtained from the actual MEG data. The goal of the M-CCA procedure is to recover dimensions in *C*. 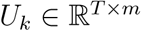 are *m* additional temporary components that are unique for each subject *k*. They are each set as a sum of 10 sinusoidal waves with their frequencies sampled randomly with the power spectrum of our MEG data. For each column or component in *U_k_*, the corresponding rows in *M_k_* are constructed by randomly selecting an “origin” dipole and the nearest *N* dipoles around the origin dipole. This is done for each subject independently. For each column or component in *C*, the same process is used to construct rows in *M_k_* for a first “seed” subject. For the rest of the 17 subjects, origin dipoles are uniformly sampled from a sphere with radius *r* centered at the origin dipole of the seed subject. Similarly, the remaining dipoles for a particular component in *C* are chosen as the nearest *N* neighbor dipoles from the origin dipole for each subject. In other words, spatial mask *M_k_* models underlying brain patterns for a particular temporal component as a cluster of dipoles, with *N* specifying the cluster size and *r* specifying the variability of their center locations across subjects. However, it is also likely that the underlying brain patterns for a particular temporal component are distributed rather than localized. To demonstrate that M-CCA is robust in both cases, in a separate simulation, we select N dipoles uniformly from the total 5124 dipoles for each temporal component without considering their coordinates.

Other than adding individual differences that create misalignment across subjects, we also consider factors that add to the misalignment across trials within a subject. Head movement blurs sensor measurement and introduces variance in downstream analyses (Stolk et al., 2013; Uutela et al., 2001). Therefore, a head movement compensation method was used as a preprocessing step on our MEG dataset before applying M-CCA (Hämäläinen & Ilmoniemi, 1994). In addition, we also investigate how head movement impacts M-CCA performance. We simulate the effect of continuous head movement by constructing a different *M_k_* for each trial produced by a shift *r*′ in the cluster centers. Shifted position in source generation mimics the effect of having a spatially fixed array but with shifted head positions.

To summarize, in the synthetic dataset, we take into account the individual differences in head geometry and sensor configuration by using the forward models obtained from experimental subjects (*F_k_*). We model the differences in structure-function mappings across subjects by modeling different parts of brain across subjects that share the same brain dynamics (*M_k_*) and by adding an additional number of unique brain dynamics that are not shared across subjects (*U_k_*). Lastly, we add different noise levels during the recording for different subjects (*w_k_*), and investigate the effect of continous head movement (*r*′). In this process, we simulate multiple trials of sensor data *S_k_* for each subject *k*. After averaging over these trials, sensor data is used as input to the M-CCA procedure.

## 3. Results

### 3.1. Synthetic Dataset

In this section, we evaluate the robustness of M-CCA against different amount of individual differences and noise levels. We measure the performance of subject alignment by how well obtained CCA components correlate across subjects. The M-CCA procedure described earlier (Figure 1) is applied to the simulated sensor data *S_k_* simultaneously for all subjects *k* = 1, 2*,…,* 18. To test for overfitting, M-CCA is applied to half of the data (even trials for each subject) as the training data to obtain the projection weights *H_k_* for each subject *k*. The same projection weights are then applied to the other half of the data (odd trials for each subject) as the testing data. Inter-subject correlations in the testing data reflect how well the CCA components truly capture the underlying data.

We generate 100 trials for each subject (50 trials as training data and 50 trials as testing data)^5^, and simulate a two-condition experiment (*T* = 200) with 100 samples per condition^6^. Signal-to-noise ratio (SNR) is set as −10 dB, which is conservative compared to typical values ( i.e. around 0 dB) approximated for real MEG datasets (Hämäläinen & Hari, 2002). for all subjects. *M_k_* selects a cluster of dipoles for each temporal component in *J_k_*, with the size of the cluster *N* set to be 200, and the variation of the cluster centers across subjects *r* set to be 0.01 m. The shared temporal components *C* used for the simulation are set to be the top 10 CCA components obtained from the actual MEG data (*n* = 10). 15 CCA components are obtained in the simulation.

Figure 2a shows that reasonable inter-subject correlations can be recovered up to the 10^th^ CCA component. There is a drop in inter-subject correlations beyond 10 CCA components, which reflects the fact that there are only 10 underlying common sources in *C* simulated in the first place. Additionally, the pattern of inter-subject correlations generalizes to testing data only up to the 10^th^ CCA component. Averaging inter-subject correlations over the top 10 CCA components (0.71), we confirmed that subject data is better aligned after the M-CCA step than before (0.34 over 306 sensors, and 0.26 over the top 10 PCA components). The low correspondence in PCA dimensions across subjects is expected, given the individual differences introduced in the simulations. The obtained CCAs are dimensions over which subjects re-align, and we can evaluate the source recovery by how well shared temporal components *C* can be expressed as a linear combination of the obtained CCAs. *Y* represents the obtained CCA components. A least-squares solution to *Q* is obtained from the system of equations *Y Q* = *C*, where *Q* transforms the obtained CCA components *Y* averaged across half of the subjects to the original common sources *C* (the other half *Y*′ is used to test for overfitting). The correlation matrix between *Y*′*Q* and *C* is calculated, with the closest correspondence for each of the 10 components in *C* being the component in *Y*′*Q* with the largest absolute correlation. The mean of these absolute correlations is 0.9791.

**Figure 2:**
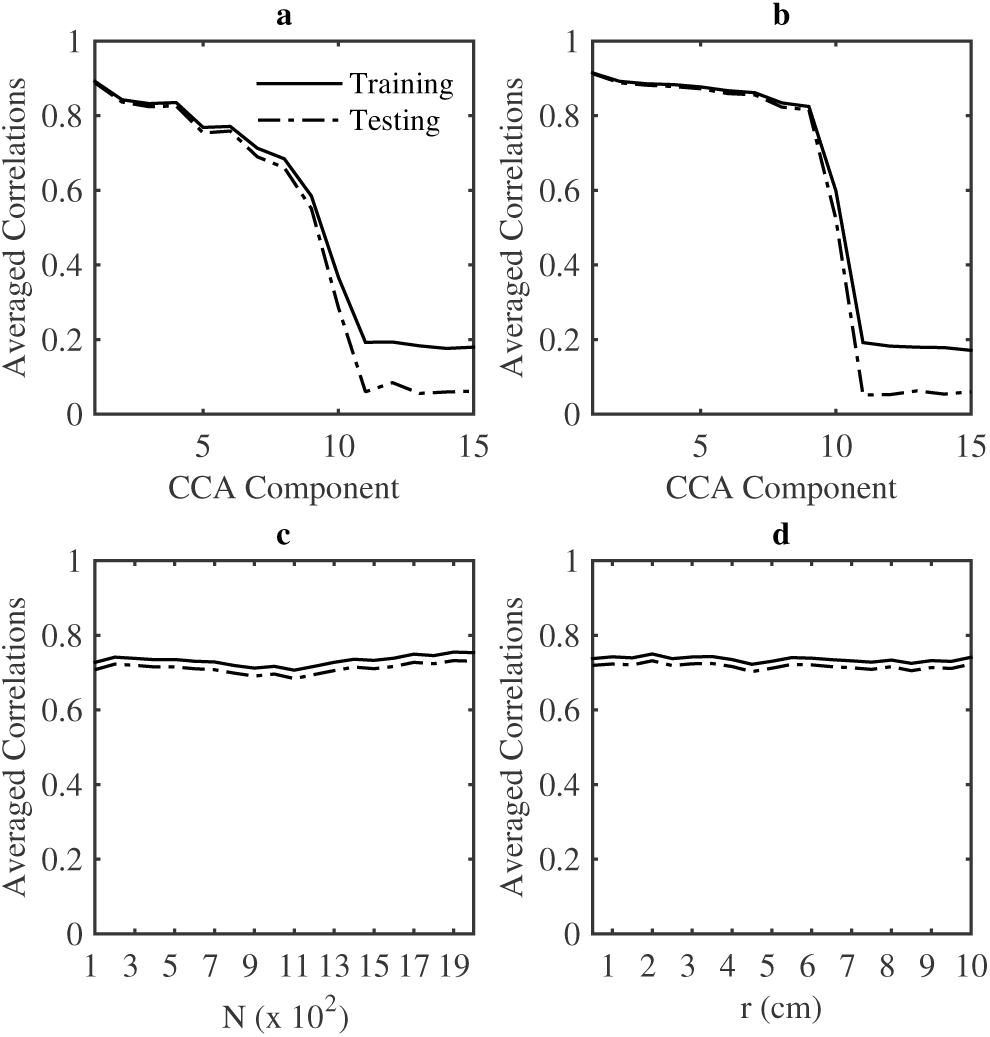
Averaged inter-subject correlations for each of the 15 CCA components while selection a cluster of dipoles of size *N* (a) or randomly select *N* dipoles (b); Inter-subject correlations averaged over the first 10 CCA components while increasing the number of dipoles *N* (c) or increasing the variation of cluster localization across subjects *r* (d). Whenever not specified, *T* = 200. SNR = −10 dB. *n* = 10. *m* = 0. *s* = 306. *N* = 200. *r* = 0.01. *r*′ = 0.01. *λ* = 0. Each data point was averaged over 10 simulations.

Next, we examine how varying different factors can have an impact on the M-CCA performance:

#### Brain patterns

(*N, r*) We examine if the M-CCA performance is affected by the pattern of dipoles (*M_k_*) that elicits a particular temporal component in *J_k_*. Figure 2a selects a cluster of dipoles for each temporal component, with the size of the cluster determined by *N* and variation of the center of the cluster across subjects determined by radius *r*. We observe that, if we instead select *N* dipoles randomly across the brain (Figure 2b), the intersubject correlations for the top 10 CCA components give similar but slightly better results than those in Figure 2a. This suggests that M-CCA is able to realign the subject data, whether the underlying brain patterns for generating the common dimensions *C* are localized (selecting a cluster of *N* dipoles) or distributed (randomly selecting *N* dipoles). For the convenience of better modeling regularity in brain sources, we continue the rest of the simulations assuming that the underlying brain patterns are clusters of dipoles. Next, we examine if the size of the cluster *N* or the radius *r* has an impact on M-CCA performance. With the rest of the parameters fixed, there is no change in the obtained inter-subject correlations varying either *N* (Figure 2c) or *r* (Figure 2c). To summarize, we have demonstrated that M-CCA performance is robust to different types of underlying brain patterns that elicit the temporal dynamics shared across subjects.

#### Signal-to-noise ratio

(*w_k_*) We vary the levels of SNR while fixing the rest of the parameters (Figure 3a). As SNR increases, the averaged inter-subject correlations of the first 10 CCA components also increases. The SNR threshold beyond which CCA components can be reasonably recovered falls into the range −30 dB to −10 dB. Until now, we have assumed that all subjects have the same SNR level. However, in reality, subjects are recorded during different experimental sessions with different noise conditions. Next, we examine if the variation in SNR level across subjects impacts M-CCA performance. We assume that SNR across subjects are sampled from a normal distribution. We consider two scenarios (two clusters of 10 points in Figure 3a). One is a low-SNR scenario where the mean of SNR across subjects is −30 dB and standard deviation ranges from 1 dB, 2 dB*,…,* 10 dB. The other represents a high-SNR scenario where the mean of SNR across subjects is 10 dB with standard deviation ranges from 1 dB, 2 dB*,…,* 10 dB. The M-CCA performance is plotted in the figure against the inter-subject correlations obtained without introducing individual differences in SNR level. Interestingly, the scattered points are clustered close together irregardless of the variation in standard deviation. It is the mean of the subject SNRs but not their variation that matters. This is the case even when the SNR mean is -30dB, where SNR from some of the subjects should fall below −30 dB given a standard deviation of 10dB.

**Figure 3:**
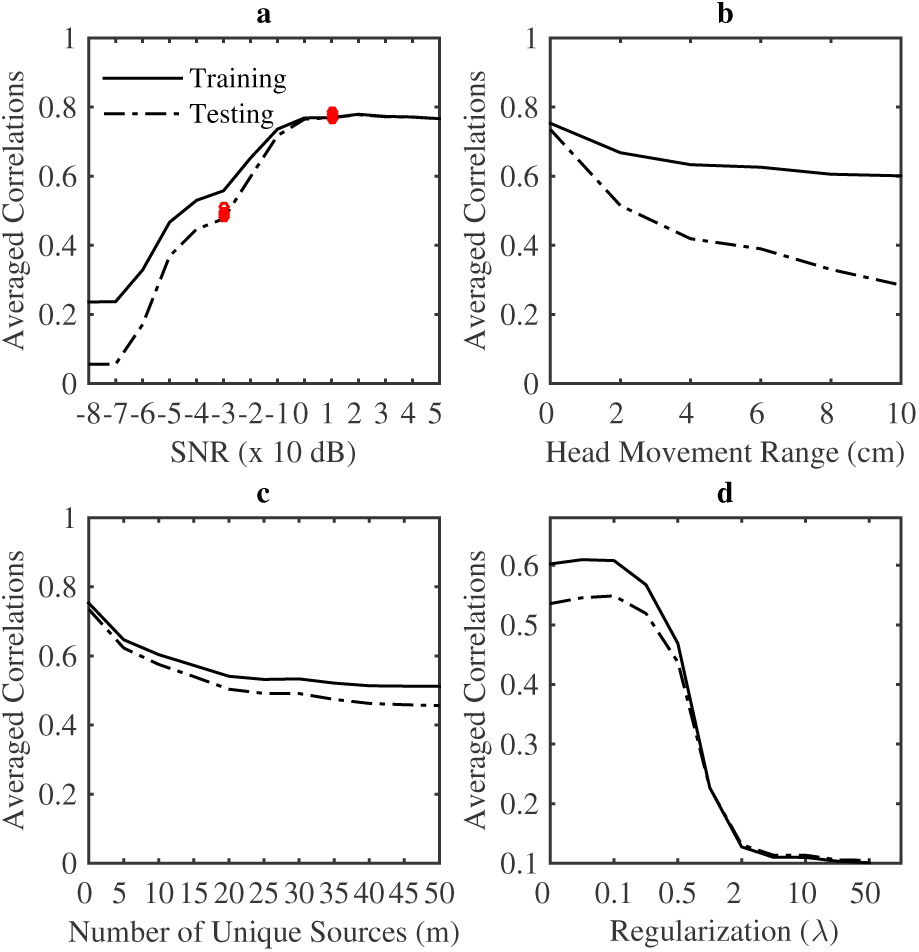
Inter-subject correlations averaged over the first 10 CCA components while increasing levels of SNR (a), the range of head movement 2*r*′ (b), the number of individual brain sources *m* (c), or when SNR = −20 dB the amount of regularization *λ* (d). Two cluster of scattered points (red) represents simulations with individual differences introduced in SNR level with *μ* = -30 dB and 10 dB for each cluster and with STD = 1 dB, 2 dB*,…,* 10 dB for each data point within each cluster. Whenever not specified, *T* = 200. SNR = −10 dB for all 18 subjects. *n* = 10. *m* = 0. *s* = 306. *N* = 200. *r* = 0.01. *r*′ = 0. *λ* = 0. Each data point was averaged over 10 simulations.

#### Head Movement

(*r*′) We generate continuous head movement across trials by introducing shifts in source generation, and examine how it can affect M-CCA performance. In particular, we vary the radius of a sphere *r*′ in which the head movement can uniformly occur (2*r*′ as the absolute range of such movement in distance). Figure 3b plots averaged inter-subject correlations while increasing 2*r*′. M-CCA performance over the testing data has reasonable inter-subject correlations up to around 1–2cm, which is comparable to challenging cases in children recordings with 0.03cm to 2.6cm as the mean shift in sensor locations (Wehner et al., 2008).

#### Addition of unique temporal components

(*U_k_*) All subjects share the same dimensions *C* in the common representational space. In addition to the shared dimensions, we vary the number of additional temporal dimensions *m* that are generated uniquely for each subject. Figure 3c shows that addition of unique temporal components for each subject decreases the M-CCA performance. But this decrease is gradual and the inter-subject correlations are reasonable up to adding 50 components. These temporal components *U_k_* are task-related brain dynamics that are locked to the stimulus across all trials in the same way as *C*. Despite the existence of such consistency, M-CCA is designed to recover only the common dimensions across subjects.

#### Addition of regularization

(*λ*) The synthetic data considers the case where subjects share similar weight maps. Under this assumption, we test if adding regularization to M-CCA improves its performance. In particular, we are interested in the situations where data has low signal-to-noise ratio (SNR = −20dB) where M-CCA may have difficulty recovering unique weight maps. In Figure 3d, over the testing data, averaged inter-subject correlations increases first over the range of 0–0.1 before it starts to decrease. Putting some emphasis to consider the similarity between sensor weight maps helps improve the inter-subject correlations of the resulted CCA components.

### 3.2. MEG Dataset

#### 3.2.1. Application of M-CCA

The M-CCA procedure described earlier (Figure 1) is applied to the MEG dataset. M-CCA is applied to the training data to obtain the projection weights *H_k_* for each subject *k*. Inter-subject correlations over the testing data are evaluated. Using 50 PCA components retained from each subject prior to M-CCA, Figure 4 shows the averaged inter-subject correlations for each of the first 20 CCA components over the training data (solid) and the testing data (dashed). The first 10 CCA components have reasonable intersubject correlations, which also generalize well over the testing data. It is reasonable to retain the first 10 CCA components for downstream analysis. In practice, there is no ground truth concerning how many dimensions there are in the common representation space. One option is to observe from which component inter-subject correlations fail to generalize to testing data. We can see, in Figure 4, that the obtained CCA components generalize well from the training data to the testing data up to around 10 CCA components. The other option is to decide a threshold of inter-subject correlations over testing data (e.g. 0.4). M-CCA ranks the obtained CCA components in terms of their importance by the inter-subject correlations, as we can observe a decreasing trend as the index of CCA component increases in Figure 4. Deciding such a threshold can be as ambiguous as deciding the amount of variance of data to keep when it comes to retaining a certain number of PCA components, which is also highly dependent on the kind of downstream analysis being carried out. If the downstream analysis is very sensitive to noise, it is advised not to include too many CCA dimensions, as subjects do not align very well in the later dimensions. If the downstream analysis is not sensitive to noise, including more CCA dimensions is useful as it keeps more information from the data.

**Figure 4:**
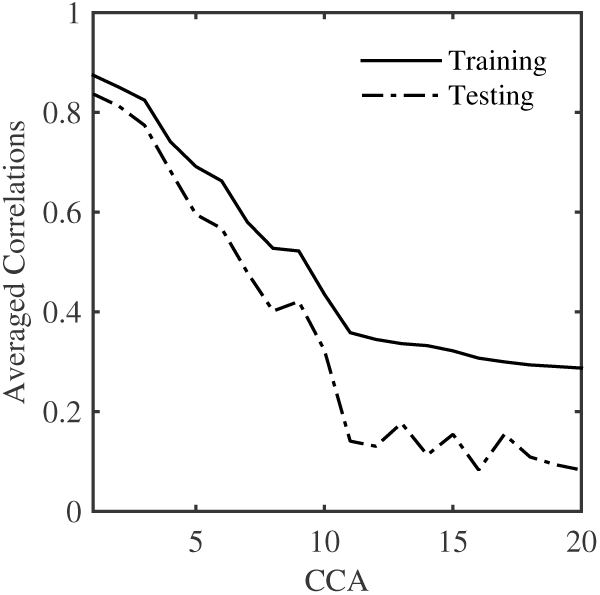
Averaged inter-subject correlations of the first 20 CCA components over training data (solid) and testing data (dashed) for the MEG dataset.

Figure 5 plots the first 10 CCA components over the 100 samples (the first 50 are stimulus-locked, and the last 50 are response-locked) for one condition. There is a considerable match between the CCA components over the training data (blue) and the testing data (magenta). Only task-relevant neural patterns that are locked to the stimulus/response and occur consistently across multiple trials will give similar temporal patterns over the training data and the testing data during the trial-averaging process. Therefore, it is unlikely that the obtained CCA components over external interference in the environment will generalize to the testing data. To further test this, we applied M-CCA to the empty room data prior to the recording of each subject^7^. High inter-subject correlations averaged over the first 10 CCA components over the training data (0.99) do not generalize over the testing data (0.07), suggesting that the top CCA components obtained in our MEG dataset do not correspond to external interference.

**Figure 5:**
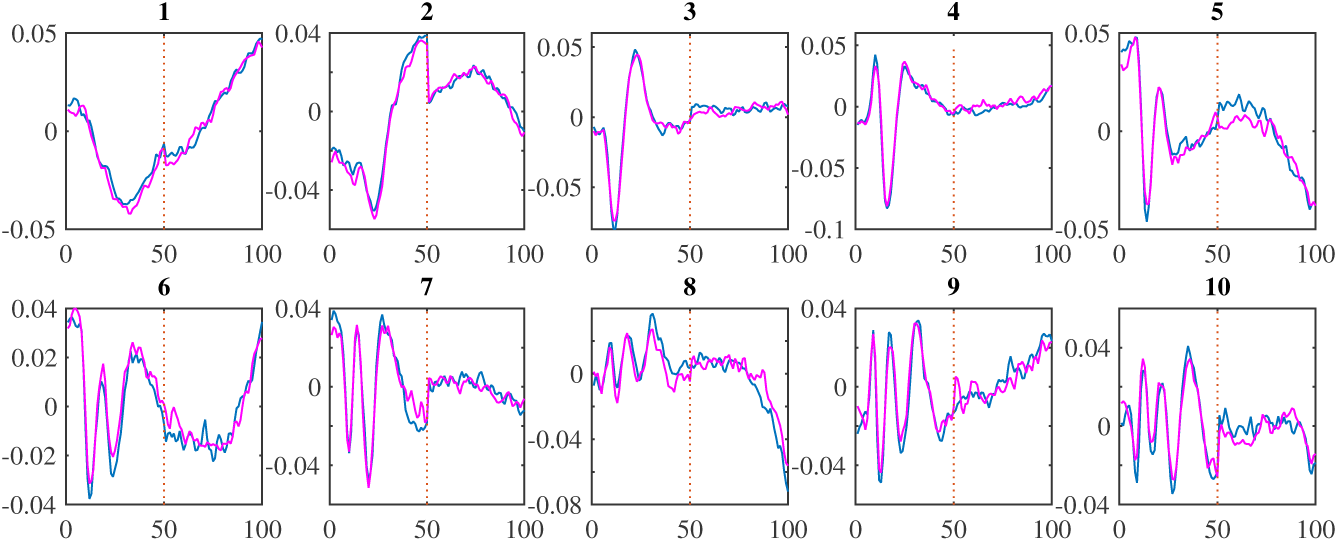
The first 10 CCA components over the first 100 samples (1 sec) averaged across all 18 subjects, with the first 50 stimulus-locked and the last 50 response-locked (separated by the red dashed line) over the training data (blue) and testing data (magenta)

The purpose of applying PCA prior to M-CCA is to reduce dimensionality and subject-specific noise. If M-CCA is applied directly to sensors, the resulting inter-subject correlation averaged over the first 10 CCA components is 0.45 for the testing data, compared to 0.59 when M-CCA is applied to 50 PCA components. Therefore, application of PCA prior to M-CCA is useful to avoid overfitting. Retaining too many PCA components may cause M-CCA to overfit, thus decreasing the inter-subject correlations, while retaining too few PCA components may lose important information. Consistent with this intuition, inter-subject correlations over the testing data first increases, then it plateaus at roughly 50–90 PCA components before decreasing. Therefore, we retained 50 PCA components prior to the M-CCA procedure over this particular MEG dataset. Using the same procedure to evaluate inter-subject correlations, the optimal number of PCA components to retain can be decided in the same way over other datasets.

#### 3.2.2. Evaluation by Inter-subject Classification

To further evaluate subject alignment using M-CCA, we compare how well we could use the transformed data to classify different experimental conditions in **inter-subject classification** and **intra-subject classification**. Inter-subject classification classifies different experimental conditions of the data from one subject given the data from other subjects, whereas intrasubject classification classifies different experimental conditions of the data from one subject given the data of the same subject. In the case when all subjects perform the task in the same way, and that data is perfectly aligned across subjects, inter-subject and intra-subject classification would yield the same classification results. Therefore, comparing inter-subject classification performance with intra-subject classification performance gives us a sense for how good subject alignment is across different methods. It is worth noting that classification accuracy also depends on intrinsic properties of the experimental condition itself: how distinct neural signals are across two different conditions, and when they are distinct. We can only meaningfully compare different classifiers/methods during the time windows where there exist distinct neural signals to classify.

We compare inter-subject classification using M-CCA with three other alternatives. The first alternative is intra-subject classification using PCA on the subjects’ sensor data, which is similar to the method used in Borst et al. (2016). This method forgoes the challenge of finding a correspondence across subjects. Second, we consider inter-subject classification using PCA on the subjects’ sensor data. Third, we consider inter-subject classification using PCA performed on source data, which has been localized using MNE and aligned based on the subjects’ anatomy. For each classification, we perform M-CCA on 100 time points for each combination of conditions excluding the condition to classify (i.e. 800 × 306 matrices instead of the 1600 × 306 matrices in Figure 1). Averaging over the dimension to be classified in M-CCA makes sure that the obtained CCA dimensions are only for maximizing the alignment across subjects, but have not learned any specific representations about the condition to classify. Classification of fan and word length are considered in the evaluation^8^.

Figure 6 and 7 show the intra-subject classification accuracy over 20 PCA components of sensor data (intraPCA: red), the inter-subject classification accuracy over 20 PCA components of the sensor data (interPCA: magenta), the inter-subject classification accuracy over 20 CCA components (interCCA: blue), and the inter-subject classification accuracy over 20 PCA components of the source data (interROI: green)^9^. Linear discriminant analysis is used for classifying data averaged over a sliding window of 10 samples, over instances each averaged over 10 trials^10^. In both Figure 6 and 7, application of M-CCA improves inter-subject classification. In fan condition (Figure 6), using CCA components has comparable classification accuracy to that of intra-subject classification, both over the stimulus-locked data and the response-locked data. Inter-subject classification accuracy using CCA components is also consistently superior to that of PCA components over the sensor data or source data. In the stimulus locked data, accuracy is above chance only after 350 ms when encoding of the words has been completed. In the response-locked data, accuracy is above chance throughout but reaches a peak about 200 ms before response generation. In word length condition (Figure 7), the period of high classifiability is early in the stimulus-locked data, during which time inter-subject classification using CCA components performs better than using PCA components over the sensor data or source data. Classification performance over the response locked data is low, given that neural signal for viewing a short word or a long word is most distinguishable during the period of time the word is being encoded (Borst et al., 2013).

**Figure 6:**
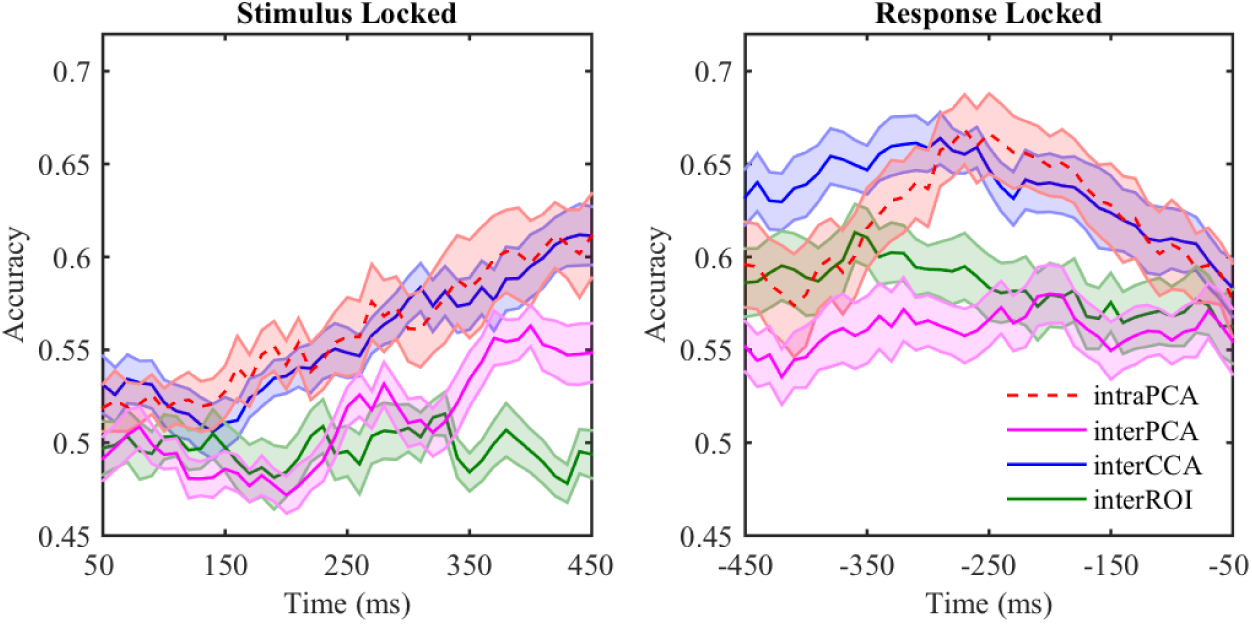
Intra-subject classification results of fan condition over PCA components of sensor data (red). Inter-subject classification results of fan condition over PCA components of sensor data (magenta), CCA components (blue), and PCA components of ROIs (red). Times on the *x* axis are relative to the stimulus (left) or the response (right). SEMs are shown in shaded error bars with *n* = 18.

**Figure 7:**
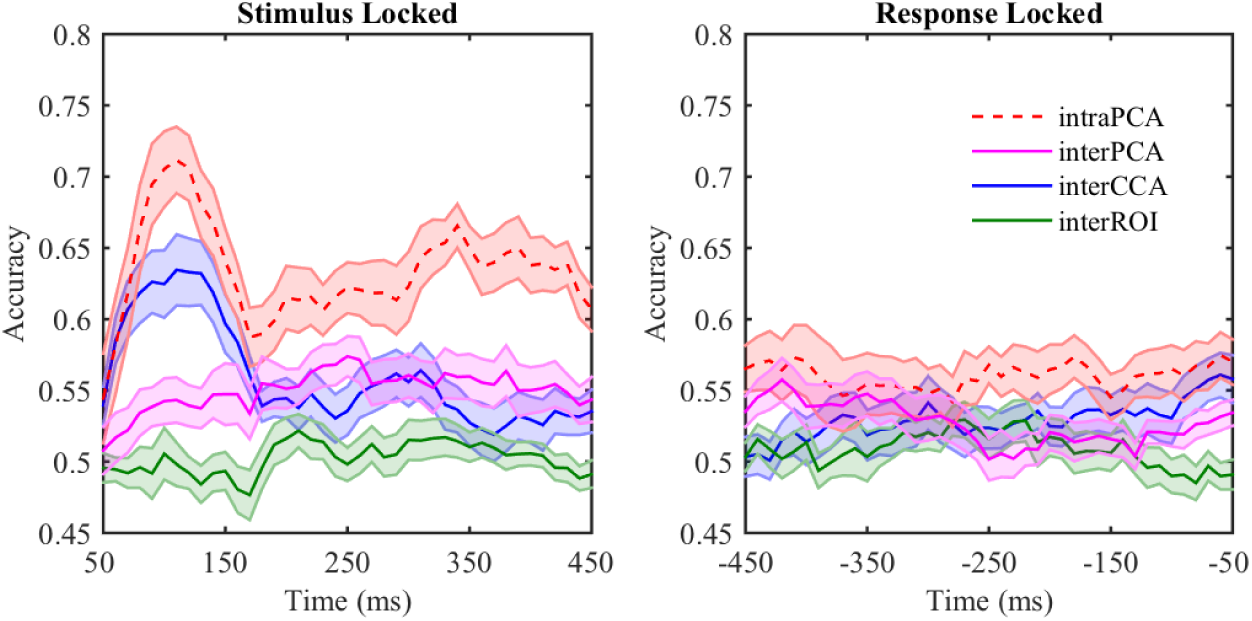
Intra-subject classification results of word length condition over PCA components of sensor data (red). Inter-subject classification results of word length condition over PCA components of sensor data (magenta), CCA components (blue), and PCA components of ROIs (green). Times on the *x* axis are relative to the stimulus (left) or the response (right). SEMs are shown in shaded error bars with *n* = 18.

#### 3.2.3. Regularization

A regularization term in the M-CCA algorithm incorporates spatial sensor information by assuming similar projection weights across different subjects. Increasing *λ* corresponds to putting more emphasis on obtaining similar weight maps, compared with obtaining more correlated CCA components across subjects. Regularization has the potential to further improve subject alignment. We evaluate this by examining the performance of inter-subject classification when *λ* is increasing. Focusing on the most classifiable period, Figure 8 shows the effect of the *λ* on inter-subject classification of fan condition during response-locked period. Each curve represents classification accuracy in a 100 ms time window centered around different times (e.g. the line marked as −100 ms refers to the inter-subject classification results during the window −150 to −50 ms). On each curve, *λ* = 0 corresponds to the same classification performance using CCA components as in Figure 6. With increasing regularization, accuracy does not decrease right away and shows a slight improvement for small *λ* values.

**Figure 8:**
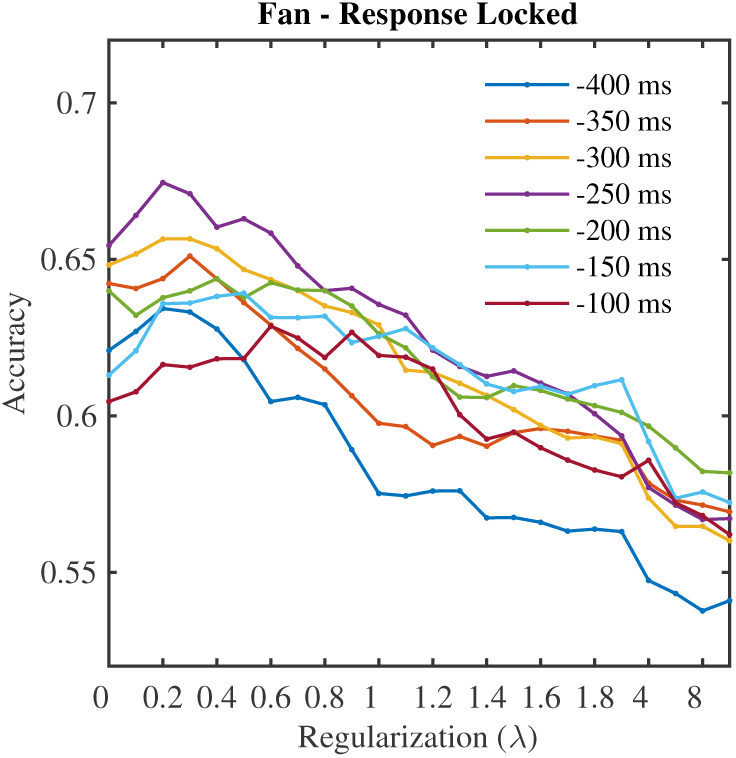
Inter-subject classification accuracies of fan condition over selected time windows of response-locked data with increasing *λ* values. Each curve represents classification accuracy in a 100 ms time window centered around different times (e.g. the line marked as −100 ms refers to the inter-subject classification results during the window −150 to −50 ms relative to the response timing).

The best *λ* value is one that is as large as possible, but does not decrease the classification accuracy much. Large *λ* values give rise to more interpretable sensor weight maps. From Figure 8 we can tell that the best range of *λ* values is around 0.2–0.4. Figure 9 shows the sensor weights that map the sensor data of the first five subjects to the first CCA component. Different columns correspond to different *λ* values ranging from 0–0.8. With increasing regularization, the change across five columns within each row/subject is minor. The difference lies in the effect of regularization on across-subject consistency of weight maps. In particular, as we move from left columns to right columns, projection weight maps across subjects become more similar. Other than the regularity of increased across-subject consistency of weight maps (which is directly imposed in the M-CCA procedure), we also observe regularity in the patterns of projection weight maps, despite the lack of any such guarantee from the regularization method itself. In particular, the sensors with higher weight values become more concentrated with increasing regularization, and they emphasize activity in symmetric lateral frontal sensors. This demonstrates that, when there are multiple solutions of weight maps at the same CCA dimension, the additional constraint in the regularization procedure may provide the most plausible and interpretable weight maps.

**Figure 9:**
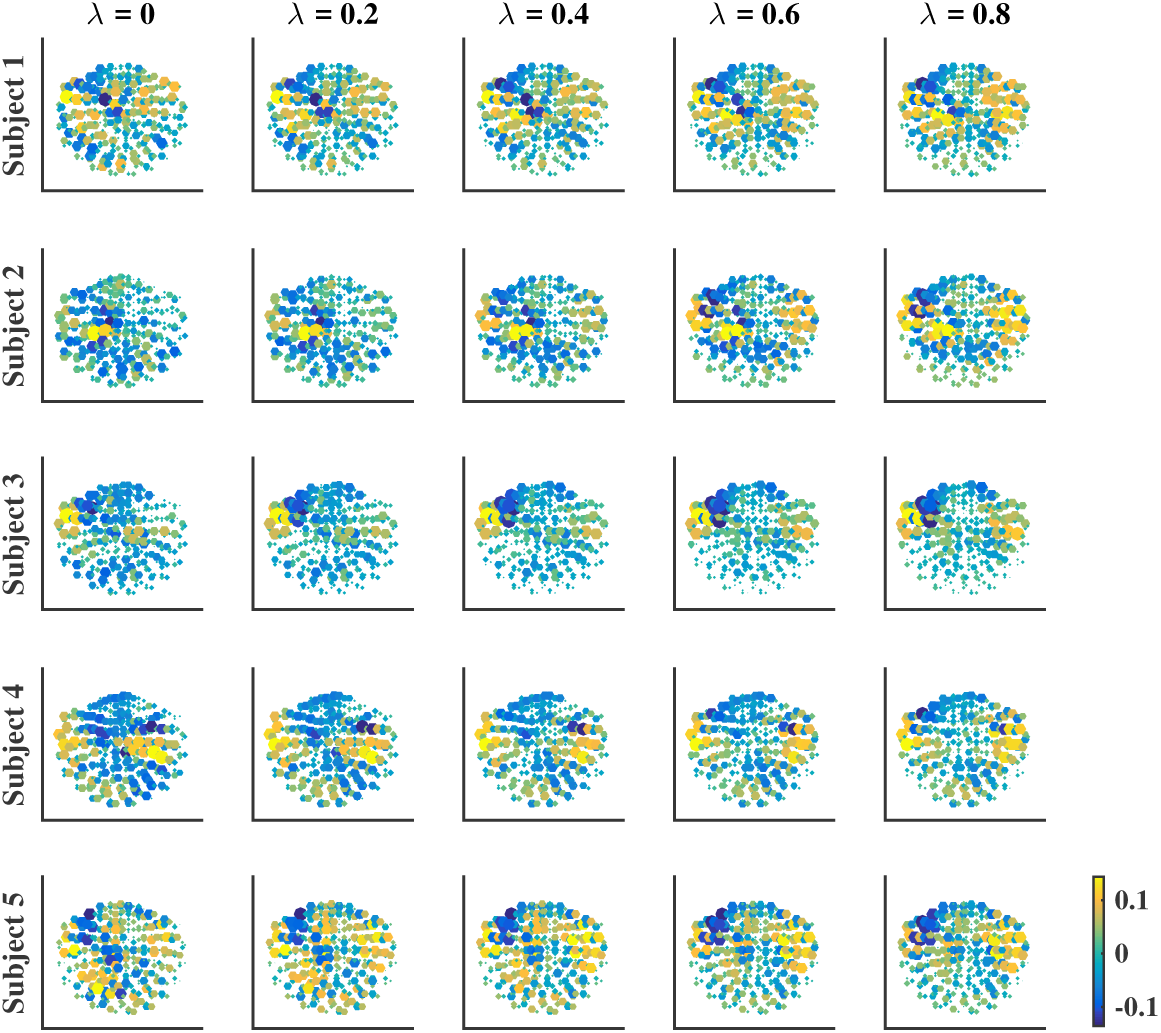
Projection weights from sensors to the first CCA component for the first five subjects with different degrees of regularization (*λ* = 0, 0.2, 0.4, 0.6, 0.8).

## 4. Discussion

We evaluated M-CCA as a method for pooling MEG data from different subjects together. It successfully produced dimensions in a common representational space over which brain activities from different subjects were well correlated, the patterns of which also generalized well over the unseen half of the data.

Subject alignment was evaluated in an inter-subject classification task, where different conditions of the data in one subject were classified based on a classifier trained on the rest of the subjects. Inter-subject classification performance using M-CCA was close to that of intra-subject classification performance over sensor data, supporting the conclusion that M-CCA succeeds in finding meaningful common dimensions. Inter-subject classification based directly on sensors never did much better than chance even though intra-subject classification based on sensors did well. This is in accordance with our knowledge that sensors do not align identically across subjects. Inter-subject classification based on source activity also performed poorly. It is likely that the quality of subject alignment using source localization was compromised by the strong assumptions that went into solving the inverse problem of source localization. Finding the inverse transformation from 306 sensors back to 5124 dipoles is a highly under-determined problem given the high dimension in dipoles (Gramfort et al., 2014). However, M-CCA is only interested in recovering the shared temporal dynamics, which are low-dimensional compared to the number of sensors. Therefore, M-CCA eliminates the need to make strong assumptions used in solving the inverse problem by directly producing the temporal dimensions that align across subjects.

We also examined the performance of M-CCA in aligning data from multiple subjects on a synthetic dataset. Realistic forward models obtained from actual experimental subjects were used to simulate different head geometries and sensor configurations across subjects. Reasonable inter-subject correlations were obtained when the SNR was larger than −10 dB, and when the continuous trial-by-trial head movement took place within the range of 1–2 cm. Other individual differences were introduced by 1) altering the spatial adjacency of the brain sources that elicit a particular shared temporal dynamics, 2) adding additional unique brain sources for each subject in the task and 3) varying the signal-to-noise ratio across subjects. M-CCA performance was robust in the presence of different types of individual differences.

In addition to the standard M-CCA algorithm, we added a regularization term to achieve better alignment and more interpretable results. M-CCA involves finding a unique mapping for each subject from the sensors to the common dimensions. The regularization term introduces a spatial constraint to impose inter-subject similarity on these sensor weight maps. This adds an appropriate constraint if the subject misalignment is the result of a small amount of spatial shifts in either sensor positions or anatomical brain regions from subject to subject. In a synthetic dataset where this kind of misalignment was present, we showed that regularization improved the recovery of the underlying sources. Over the real MEG dataset, adding the regularization term also improved the inter-subject classification performance and produced more consistent and interpretable sensor weight maps across subjects.

In this paper, we outlined the pipeline and evaluated the suitability of using M-CCA to align subjects in MEG data. The eighteen subjects were recorded in the same MEG environment. However, given the insensitivity of M-CCA to variation of noise conditions across different recordings (demonstrated in the synthetic dataset), M-CCA can be applied when subjects are recorded under different sites or even under different MEG systems, provided that we can still assume the existence of a linear transformation from source activity to sensor activity. In addition, the use of M-CCA in pooling data from different subjects is not restricted to only one neural imaging modality. We demonstrated in this study how M-CCA can be applied to align subjects in MEG that comes in at a fine temporal grain size. In this aspect, EEG (Electroencephalogram) and ECoG (Electrocorticography) datasets share very similar temporal characteristics as MEG, and can utilize M-CCA to pool subjects together in the same way. Lastly, it is possible to pool subjects recorded from different studies of the same task, as long as there are corresponding time points across subjects and that brain responses over that period of the time are considered consistent across subjects.

## 5. Acknowledgements

This research was supported by the National Science Foundation grant 1420009 to J.R.A., the James S. McDonnell Foundation Scholar Award 220020162 to J.R.A., the Office of Naval Research grant N00014-15-1-2151 to J.R.A., the National Institute of Mental Health grant 064537 to R.E.K., and the Netherlands Organization for Scientific Research Veni grant 451-15-040 to J.P.B.. We thank Matthew M. Walsh and Aryn Pyke for their helpful comments on a draft of the paper. The Authors declare that there is no conflict of interest.

## Footnotes

1 Note that we use ‘sensor’ to indicate the magnetic coils outside of the head and ‘source’ to indicate the origin of the measured signal on the cortex.

2 Typically the standard MNI305 brain is used.

3 Maximizing the consistency across different subjects has been previously used to align fMRI data in human ventral temporal cortex (Haxby et al., 2011). Our proposed method, in addition, simultaneously reduces dimensionality as the dimensions in the common representational space are ranked by how well subjects align.

4 Based on Nyquist sampling criterion, 100 Hz sampling rate is sufficient to capture the neural signal filtered at 0.5-50 Hz.

5 It is comparable to 54.7 trials per condition per subject in the real MEG dataset.

6 There are 16 experimental conditions available in the real MEG dataset, but here we simulate a more general case with only two experimental conditions.

7 Non-overlapping segments of data of 100 samples are taken as trials when applying M-CCA. On average there are 272 trials available in the empty room data per subject.

8 Probe type condition is not included due to inferior intra-subject classification performance. Trials responded by left hand and right hand are so different that they have to be distinguished in the CCA step in order to have reasonable functional alignment. As a result, left and right hand condition cannot be taken as a condition to classify.

9 Classification performance does not improve further adding more components beyond 20 for both PCA components over sensors/ROIs and CCA components.

10 None of the classifiers have satisfactory performance over single trials, thus an effective comparison among the classifiers is not possible.

## References

Anderson, J. R., Zhang, Q., Borst, J. P., & Walsh, M. M. (2016). The discovery of processing stages: Extension of sternberg’s method. Psychological review, 123, 481–509.

Borga, M. (1998). Learning Multidimensional Signal Processing. Ph.D. thesis Linkoping University.

Borst, J. P., Ghuman, A. S., & Anderson, J. R. (2016). Tracking cognitive processing stages with meg: A spatio-temporal model of associative recognition in the brain. NeuroImage, 141, 416–430.

Borst, J. P., Schneider, D. W., Walsh, M. M., & Anderson, J. R. (2013). Stages of processing in associative recognition: Evidence from behavior, eeg, and classification. Journal of Cognitive Neuroscience, 25, 2151–2166. URL: http://dx.doi.org/10.1162/jocn_a_00457. doi:10.1162/jocn_a_00457.

Correa, N. M., Adali, T., Li, Y.-O., & Calhoun, V. D. (2010). Canonical correlation analysis for data fusion and group inferences: Examining applications of medical imaging data. IEEE signal processing magazine, 27, 39–50.

Dale, A. M., Fischl, B., & Sereno, M. I. (1999). Cortical surface-based analysis: I. segmentation and surface reconstruction. NeuroImage, 9, 179–194.

Fischl, B. (2012). Freesurfer. NeuroImage, 62, 774–781.

Gramfort, A., Luessi, M., Larson, E., Engemann, D. A., Strohmeier, D., Brodbeck, C., Parkkonen, L., & Hämäläinen, M. S. (2014). MNE software for processing MEG and EEG data. NeuroImage, 86, 446–460.

Gramfort, A., Papadopoulo, T., Olivi, E., & Clerc, M. (2010). Openmeeg: opensource software for quasistatic bioelectromagnetics. BioMedical Engineering OnLine, 9, 45. URL: http://dx.doi.org/10.1186/1475-925X-9-45. doi:10.1186/1475-925x-9-45.

Hämäläinen, M., & Hari, R. (2002). Magnetoencephalographic (meg) characterization of dynamic brain activation, .

Hämäläinen, M., Hari, R., Ilmoniemi, R. J., Knuutila, J., & Lounasmaa, O. V. (1993). Magnetoencephalography—theory, instrumentation, and applications to noninvasive studies of the working human brain. Reviews of Modern Physics, 65, 413–497. URL: http://dx.doi.org/10.1103/RevModPhys.65.413. doi:10.1103/revmodphys.65.413.

Hamalainen, M., & Sarvas, J. (1989). Realistic conductivity geometry model of the human head for interpretation of neuromagnetic data. IEEE Transactions on Biomedical Engineering, 36, 165–171. URL: http://dx.doi.org/10.1109/10.16463. doi:10.1109/10.16463.

Hämäläinen, M. S., & Ilmoniemi, R. J. (1994). Interpreting magnetic fields of the brain: minimum norm estimates. Med. Biol. Eng. Comput., 32, 35–42. doi:10.1007/bf02512476.

Haxby, J. V., Guntupalli, J. S., Connolly, A. C., Halchenko, Y. O., Conroy, B. R., Gobbini, M. I., Hanke, M., & Ramadge, P. J. (2011). A common, high-dimensional model of the representational space in human ventral temporal cortex. Neuron, 72, 404–416. URL: http://dx.doi.org/10.1016/j.neuron.2011.08.026. doi:10.1016/j.neuron.2011.08.026.

Kettenring, J. R. (1971). Canonical analysis of several sets of variables. Biometrika, 58, 433–451.

Larson, E., & Taulu, S. (2016). The importance of properly compensating for head movements during meg acquisition across different age groups. Brain Topography, 30, 172–181. URL: http://dx.doi.org/10.1007/s10548-016-0523-1. doi:10.1007/s10548-016-0523-1.

Li, Y.-O., AdalÄś, T., Wang, W., & Calhoun, V. D. (2009). Joint blind source separation by Multi-set Canonical Correlation analysis. IEEE transactions on signal processing: a publication of the IEEE Signal Processing Society, 57, 3918–3929.

Li, Y.-O., Eichele, T., Calhoun, V. D., & Adali, T. (2012). Group study of simulated driving fMRI data by multiset canonical correlation analysis. Journal of Signal Processing Systems, 68, 31–48.

Mosher, J. C., Leahy, R. M., & Lewis, P. S. (1999). Eeg and meg: forward solutions for inverse methods. IEEE Transactions on Biomedical Engineering, 46, 245–259.

Nenonen, J., Nurminen, J., Kičić, D., Bikmullina, R., Lioumis, P., Jousmäki, V., Taulu, S., Parkkonen, L., Putaala, M., & Kähkönen, S. (2012). Validation of head movement correction and spatiotemporal signal space separation in magnetoencephalography. Clinical Neurophysiology, 123, 2180–2191. URL: http://dx.doi.org/10.1016/j.clinph.2012.03.080. doi:10.1016/j.clinph.2012.03.080.

Norman, K. A., Polyn, S. M., Detre, G. J., & Haxby, J. V. (2006). Beyond mind-reading: multi-voxel pattern analysis of fmri data. Trends in Cognitive Sciences, 10, 424–430.

Oostendorp, T., & van Oosterom, A. (1989). Source parameter estimation in inhomogeneous volume conductors of arbitrary shape. IEEE Transactions on Biomedical Engineering, 36, 382–391. URL: http://dx.doi.org/10.1109/10.19859. doi:10.1109/10.19859.

Oostenveld, R., Fries, P., Maris, E., & Schoffelen, J.-M. (2011). Fieldtrip: Open source software for advanced analysis of meg, eeg, and invasive electrophysiological data. Computational Intelligence and Neuroscience, 2011, 1–9. URL: http://dx.doi.org/10.1155/2011/156869. doi:10.1155/2011/156869.

Picton, T., Bentin, S., Berg, P., Donchin, E., Hillyard, S., Johnson, R., Miller, G., Ritter, W., Ruchkin, D., Rugg, M., & Taylor, M. (2000). Guidelines for using human event-related potentials to study cognition: Recording standards and publication criteria. Psychophysiology, 37, 127–152.

Rustandi, I., A., J. M., & Mitchell, T. M. (2009). Integrating multiple-study multiple-subject fMRI datasets using canonical correlation analysis.

Scherg, M., & Von Cramon, D. (1985). Two bilateral sources of the late aep as identified by a spatio-temporal dipole model. Electroencephalography and Clinical Neurophysiology/Evoked Potentials Section, 62, 32–44. URL: http://dx.doi.org/10.1016/0168-5597(85)90033-4. doi:10.1016/0168-5597(85)90033-4.

Stolk, A., Todorovic, A., Schoffelen, J.-M., & Oostenveld, R. (2013). Online and offline tools for head movement compensation in meg. NeuroImage, 68, 39–48. URL: http://dx.doi.org/10.1016/j.neuroimage.2012.11.047. doi:10.1016/j.neuroimage.2012.11.047.

Uutela, K., Taulu, S., & Hämäläinen, M. (2001). Detecting and correcting for head movements in neuromagnetic measurements. NeuroImage, 14, 1424–1431. URL: http://dx.doi.org/10.1006/nimg.2001.0915. doi:10.1006/nimg.2001.0915.

Vía, J., Santamaría, I., & Pérez, J. (2005). Canonical correlation analysis (CCA) algorithms for multiple data sets: Application to blind simo equalization.

Vía, J., Santamaría, I., & Pérez, J. (2007). A learning algorithm for adaptive canonical correlation analysis of several data sets. Neural Networks, 20, 139–152.

Wehner, D. T., Hämäläinen, M. S., Mody, M., & Ahlfors, S. P. (2008). Head movements of children in meg: Quantification, effects on source estimation, and compensation. NeuroImage, 40, 541–550. URL: http://dx.doi.org/10.1016/j.neuroimage.2007.12.026. doi:10.1016/j.neuroimage.2007.12.026.

